# New distribution records and range extensions of mosquitoes in British Columbia and the Yukon Territory

**DOI:** 10.1101/2020.01.24.919233

**Authors:** Daniel AH. Peach, Lisa M. Poirier

## Abstract

We report the first records of *Aedes euedes* Howard, Dyar, and Knab, and *Coquillettidia perturbans* (Walker) from Canada’s Yukon Territory, and the first record of *Ae. decticus* Howard, Dyar, and Knab from British Columbia. We also report range extensions in northern BC for the western treehole mosquito, *Aedes sierrensis* (Ludlow), the common house mosquito, *Culex pipiens* L., and the cool weather mosquito *Culiseta incidens* (Thomson).

## Introduction

Large-scale mosquito trapping for West Nile virus (WNV) surveillance has yielded several additions to the mosquito fauna of BC (Peach 2018b). This effort was logically focused on the south of the province, where most of the human population is concentrated and where WNV is most likely to occur. In contrast, northern BC and the Yukon Territory have experienced much lower survey effort, with bioblitzes and individual collecting efforts largely responsible for advances in our knowledge of the mosquito fauna in these areas.

50 mosquito species are currently known from British Columbia (Peach 2018b), and 31 from the Yukon Territory (Belton and Belton 1990; Peach 2017, 2018a). Here we present new distribution records and range extensions from recent mosquito collecting efforts, paired with supporting historical records where available and relevant. We highlight the first records of *Coquillettidia perturbans* (Walker), and *Aedes euedes* Howard, Dyar, and Knab, from Canada’s Yukon Territory, and the first record of *Ae. decticus* from British Columbia. While some of these records surely represent species that were historically present and simply went undetected, others may represent more recent extensions of northern range limits brought on by changing climate, anthropogenic dispersal, or an increase of synanthropic habitat availability.

## Results

### *Aedes decticus* Howard, Dyar, and Knab

*Aedes decticus* Howard, Dyar, and Knab is an uncommon but widely distributed mosquito (Wood et al. 1979). It lacks transverse basal bands of pale scales on its abdominal tergites and has black scales on its vertex and postpronotum (Wood *et al*. 1979). Little is known of its life history, but it has been reported in sphagnum bogs and other acidic woodland bogs and pools (Mullen 1971). *Ae. decticus* has been reported from New England, regions surrounding the Great Lakes, Labrador, northern Manitoba, and Alaska (Wood *et al*. 1979). *Ae. decticus* has been previously recorded from the Yukon Territory (Belton and Belton 1990), but this has been overlooked in recent literature (Darsie and Ward 2005). Adult *Ae. decticus* lack a transverse basal band of pale scales on the bases of abdominal tergites and have a mix of yellow and dark scales present on the vertex and postpronotum (Wood *et al*. 1979; Darsie and Ward 2005). *Ae. decticus* will take blood meals from humans (Smith 1952), but little is known about what other sources of vertebrate blood it will take.

One adult female *Ae. decticus* was collected while attempting to feed on DP at Wye Lake, YT, on July 12, 2019, and another while attempting the same at a bog just north of the town of Watson Lake on July 13, 2019 (Table 1). Another adult female *Ae. decticus* was collected while attempting to feed on DP at Morley Lake, BC, about 100 m south of the BC/Yukon border on July 15, 2019.

**Table 1:**
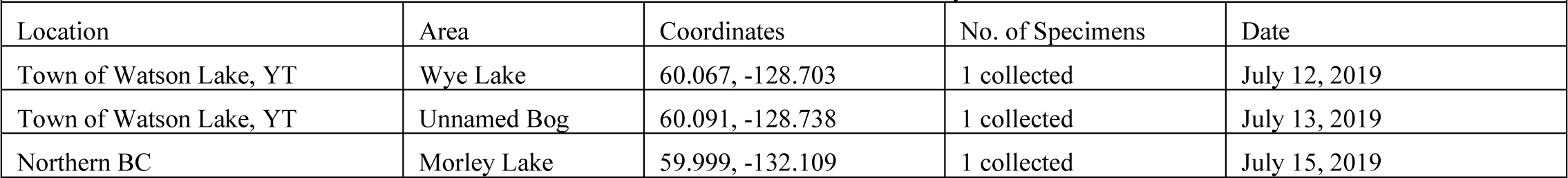
Collection records of *Ae. decticus* from British Columbia and the Yukon Territory.

| Location | Area | Coordinates | No. of Specimens | Date |
| --- | --- | --- | --- | --- |
| Town of Watson Lake, YT | Wye Lake | 60.067, -128.703 | 1 collected | July 12, 2019 |
| Town of Watson Lake, YT | Unnamed Bog | 60.091, -128.738 | 1 collected | July 13, 2019 |
| Northern BC | Morley Lake | 59.999, -132.109 | 1 collected | July 15, 2019 |

*Ae. decticus* is undoubtedly present on both sides of the border at the Morley Lake location. While the extent of its range is unknown, the extensive spruce forest and boggy conditions present in northern BC and the Yukon likely provide many localized areas of suitable habitat for this species, and such conditions may extend from Alaska to Labrador.

### Aedes euedes Howard, Dyar, and Knab

*Aedes eueudes* Howard, Dyar, and Knab is a large, uncommon mosquito with bands of pale scales on the tarsi and scattered pale scales on the proboscis, cerci, and beyond the pale basal bands of the abdominal tergites (Wood *et al*. 1979; Belton 1983; Darsie and Ward 2005). It lacks lower mesepimeral setae and possesses reddish brown scales on the scutum (Wood *et al*. 1979; Belton 1983; Darsie and Ward 2005). It is thought to complete only one generation per year and to breed in permanent or semi-permanent pools in a variety of habitats including woodland, marshes, and grassy areas (Wood *et al*. 1979; Westwood *et al*. 1983).

*Ae. euedes* has been found in BC, Alberta, the Northwest Territories, and Alaska (Wood *et al*. 1979; Belton 1983; Darsie and Ward 2005; Peach 2018b). *Ae. euedes* was thought to likely be present but undetected in the Yukon Territory (Belton and Belton 1990). In summer 2019 adult females were collected while trying to bite DP in an open area near wastewater treatment lagoons outside of Whitehorse, YT, in forest on the margins of the Liard River, and along the margins of Seaforth Creek where it crosses the Alaska Highway (Table 2). This last area was a grassy riparian habitat that contained numerous willows and small marshy areas with surrounding northern forest.

**Table 2:**
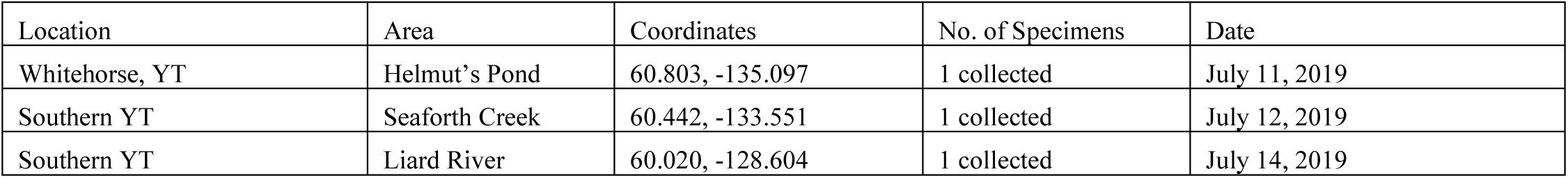
Collection records of *Ae. euedes* from the Yukon Territory.

These collections represent the first confirmed records of *Ae. euedes* from the Yukon Territory. This species is likely present at low numbers in suitable habitat throughout the Yukon Territory, as it has been found as far north as the coast of the Arctic Ocean in the neighbouring Northwest Territories (Wood *et al*. 1979).

### Aedes sierrensis (Ludlow)

The western treehole mosquito, *Aedes sierrensis* (Ludlow), is a multivoltine species that breeds in water-filled holes in trees and can also occasionally be found in water-filled man-made containers rich in plant debris and leaves. Adults are black with striking bands of silvery-white scales on the legs and patterns of the same on the scutum.

*Ae. sierrensis* is an aggressive day-biter that takes blood from warm-blooded animals, including humans (Peyton 1956), and is a vector of dog heartworm, *Dirofilaria immitis* (Belton 1983). *Aedes sierrensis* is found over a broad portion of western North America, from California to northern BC (Belton 1983; Darsie and Ward 2005), wherever mature trees with suitable larval habitat are found. However, it was not previously known to occur north of Terrace or on the Islands of BC’s north coast. Historical specimens in the Royal BC Museum indicate the presence of *Ae. sierrensis* in the vicinity of Haida Gwaii, and an additional specimen collected in August 2019 by Chris Stinson in Queen Charlotte City confirms the presence of this species in the area (Table 3). The range of this species may also extend northward into the Alexander Archipelago.

**Table 3:**
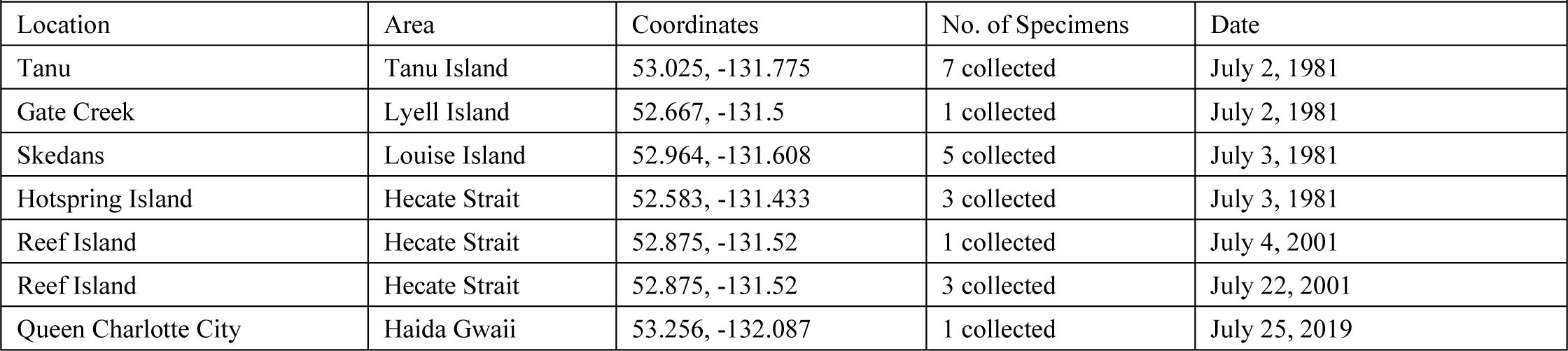
Collection records of *Ae. sierrensis* from the North Coast of British Columbia.

| Location | Area | Coordinates | No. of Specimens | Date |
| --- | --- | --- | --- | --- |
| Tanu | Tanu Island | 53.025, -131.775 | 7 collected | July 2, 1981 |
| Gate Creek | Lyell Island | 52.667, -131.5 | 1 collected | July 2, 1981 |
| Skedans | Louise Island | 52.964, -131.608 | 5 collected | July 3, 1981 |
| Hotspring Island | Hecate Strait | 52.583, -131.433 | 3 collected | July 3, 1981 |
| Reef Island | Hecate Strait | 52.875, -131.52 | 1 collected | July 4, 2001 |
| Reef Island | Hecate Strait | 52.875, -131.52 | 3 collected | July 22, 2001 |
| Queen Charlotte City | Haida Gwaii | 53.256, -132.087 | 1 collected | July 25, 2019 |

### Culex pipiens L

The northern house mosquito, *Culex pipiens* L., is a multivoltine mosquito native to Eurasia that breeds in stagnant and polluted standing water including ditches, sewage lagoons, and water-filled containers rich in organic material (Wood *et al*. 1979; Belton 1983). Larvae have an elongated siphon and are often found feeding at the water’s surface. Adult *Cx. pipiens* are brown with bands of pale scales on the bases of abdominal tergites. This species overwinters as nulliparous adult females that take shelter in locations such as sheds, rodent burrows, tree bark, and rock piles until the following spring when they emerge to feed.

*Cx. pipiens* blood-feed primarily from birds, though they will feed on humans as well (*Wood et al*. 1979). These feeding habits make *Cx. pipiens* important vectors of West Nile virus (Hamer et al. 2008). *Cx. pipiens* have also been observed feeding on a variety of nectar sources, including yarrow, *Achillea millefolium*, and common tansy, *Tanacetum vulgare* (Peach and Gries 2019), which they pollinate (Peach and Gries 2016).

Previously, *Cx. pipiens* had not been reported north of southern BC (Belton 1983; Darsie and Ward 2005); however, in 2004, two specimens were captured in a CDC blacklight trap in Valemount, BC by T. Brown and two more were captured at a sewage lagoon in Prince George, BC. Follow-up trapping conducted by LP in 2019, 15 years later, at the Prince George site resulted in the capture of an additional 8 adult female *Cx. pipiens* (Table 4). Suspected *Cx. pipiens* were observed by DP in Kitimat during September 2019 but evaded capture.

**Table 4:** Northern records of *Cx. pipiens*.

| **Table 4:** Northern records of *Cx. pipiens*. |  |  |  |  |
| --- | --- | --- | --- | --- |
| Location | Area | Coordinates | No. of Specimens | Date |
| Valemount | Cranberry Marsh/Starratt Wildlife Management Area | 52.816, -119.268 | 1 collected | Aug. 19, 2004 |
| Valemount | Cranberry Marsh/Starratt Wildlife Management Area | 52.819, -119.259 | 1 collected | Sept. 9, 2004 |
| Prince George | Danson Lagoon | 53.831, -122.735 | 2 collected | Aug. 11, 2004 |
| Prince George | Danson Lagoon | 53.831, -122.735 | 8 collected | Aug. 27, 2019 |

It is likely that cold northern winters restrict the northern limits of *Cx. pipiens* distribution; however, as the species will overwinter in heated manmade structures it is possible that it can be found in human settlements with suitable breeding conditions much farther north than would be expected due to the presence of warm outbuildings and human transportation of equipment or other materials that may contain adults or eggs.

### Culiseta incidens (Thomson)

The cool weather mosquito, *Culiseta incidens* (Thomson), is BC’s most common and widespread mosquito species (Belton 1983). *Cs. incidens* is a large mosquito with aggregations of dark scales on the wings and very narrow bands of pale scales on the tarsi. This mosquito breeds in artificial containers, ditches, polluted water, storm drains, woodland pools, and coastal and riverine rock pools. Females overwinter in warm, dry locations such as sheds, under tree bark, rock piles, or rodent burrows.

*Cs. incidens* is the most common and widespread mosquito species in BC and is found in almost every part of the province. However, it was not previously known to occur from Haida Gwaii and other islands on BC’s North Coast. Historical specimens in the Royal BC Museum indicate the presence of *Cs. incidens* on Haida Gwaii and other islands on BC’s North Coast, and an additional specimen collected in July 2019 by Megan Willie confirms the presence of this species in the area (Table 5). The range of this species may also extend northward into the Alexander Archipelago.

**Table 5:** Collection records of *Cs. Incidens* from the North Coast of British Columbia

|  |  |  |  |  |
| --- | --- | --- | --- | --- |
| Location | Area | Coordinates | No. of Specimens | Date |
| Ross Island | Hecate Strait | 52.163, -131.121 | 2 collected | June 30, 1981 |
| Gate Creek | Lyell Island, Haida Gwaii | 52.665, -131.466 | 3 collected | July 2, 1981 |
| Tanu Island | Hecate Strait | 52.766, -131.617 | 1 collected | July 2, 1981 |
| Petrel Islet | Hippa Island | 53.551, -133.011 | 1 collected | July 24, 2019 |

### Coquillettidia perturbans (Walker)

The cattail mosquito, *Coquillettidia perturbans* (Walker), is a univoltine mosquito that breeds in shallow bodies of water such as swamps, marshes, or shallow lakes, with abundant emergent vegetation and a layer of soft mud or peat at the bottom (Carpenter and LaCasse 1955; Wood *et al*. 1979; Belton 1983). Larvae have a highly-modified siphon to attach to and obtain oxygen from the roots of emergent aquatic plants (Wood et al. 1979), and this species overwinters anchored to these roots, submerged in mud (Carpenter and LaCasse 1955; Wood *et al*. 1979). *Cq. perturbans* adults possess bands of pale scales on the tarsi, a band of pale scales in the middle of the tibia, and a mix of large, triangular, black and white scales on the wing veins (Carpenter and LaCasse 1955; Wood *et al*. 1979; Belton 1983). Adults have been reported to feed on nectar from flowers of goldenrod (*Solidago spp*.), yarrow (*Asclepias millefolium*), milkweed (*Asclepias spp*.), dogbane (*Apocynum spp*.), and more (Sandholm and Price 1962; Grimstad and DeFoliart 1974).

*Cq. perturbans* is aggressive, mammalophilic, and a vector of West Nile virus (WNV) and eastern equine encephalitis virus (EEEV) (Turell et al. 2005). It has often been described as having a southerly distribution (Wood *et al*. 1979; Belton 1983); however, it has previously been recorded as far North as Fort Nelson, British Columbia (Poirier and Berry 2011).

During the July 2019 Yukon Bioblitz DP collected several adult *Cq. perturbans* females attempting to blood-feed at two locations in the southern Yukon Territory and one location in northern British Columbia (Table 6).

**Table 6:** Collection records of *Cq. perturbans* from British Columbia and the Yukon Territory.

|  |  |  |  |  |
| --- | --- | --- | --- | --- |
| Location | Area | Coordinates | No. of Specimens | Date |
| Wye Lake | Town of Watson Lake, YT | 60.065, -128.701 | 2 collected | July 12, 2019 |
| Albert Creek bird banding station | Upper Liard, YT | 60.062, -128.917 | 3 observed, 2 collected | July 13, 2019 |
| Coal River Lodge | Coal River, BC | 59.657, -126.956 | 2 collected | July 14, 2019 |

The collections from Upper Liard and Watson Lake represent the first records of this species and genus in the Yukon Territory and are more than 100 km north and 300 km west of Fort Nelson, previously the most northerly known collection record for *Cq. perturbans* (Poirier and Berry 2011). Both locations were shallow bodies of water with ample emergent vegetation, including *Nuphar variegate, Equisetum fluviatile*, and *Persicaria amphibia*, and were within the boreal cordillera ecozone (Canadian Council on Ecological Areas 1996).

The collection site at Coal River, BC was in forest beside the Alaska Highway, but many oxbow lakes are present in the area and we believe our specimens may have originated from these.

## Acknowledgements

We thank the Regional District of Fraser-Fort George for funding of the 2004 mosquito trapping program, the Royal BC Museum and the Beaty Biodiversity Museum for access to their specimens, as well as Taryn Brown, Chris Stinson, Megan Willie, and all of those who have collected mosquito specimens referred to in this manuscript. We also thank the organizers of the Yukon Bioblitz for this fantastic event.

